# Hubness and local scaling in paired T cell receptor distance spaces

**DOI:** 10.64898/2026.10.13.758726

**Authors:** Kilian Maire, Wenning Zheng

## Abstract

Nearest-neighbour retrieval in T cell receptor distance spaces is uneven: some receptors enter many neighbour lists while others enter none. We quantified this hubness in paired variable-domain distances and evaluated local scaling as a post-processing correction. Across ITRAP-filtered dextramers, VDJdb and McPAS-TCR, the skewness of the ten-neighbour occurrence count was 1.26 to 1.50. Local scaling reduced it to 0.13 to 0.31.

Under a diagnostic restriction requiring different V families on both chains, macro retrieval gain increased by 0.1418 on VDJdb (95% hierarchical bootstrap interval 0.0079 to 0.3016). The same untuned correction gave a gain of 0.1185 on the held-out IMMREP25 corpus (0.0040 to 0.2477). On IMMREP22, mean micro-AUC increased from 0.8091 to 0.8394, with improvement on 14 of 17 epitopes (paired Wilcoxon p = 0.0026). Gains for CDR3 edit distance and SCEPTR had the same direction but were not significant separately.

The benefit has narrow boundaries. CDR3 length alone recovered approximately one-third to three-fifths of it. Eight corrections failed to give reproducible gains on deduplicated beta-only receptors in the two principal corpora. Unrestricted retrieval did not yield a gain surviving the declared multiplicity corrections. Directly penalizing neighbour occurrence was harmful, and clustering results depended on partition granularity. Hubness reduction and specificity retrieval are therefore distinct outcomes. These findings support reporting hubness alongside retrieval accuracy and testing local scaling within paired-domain evaluations; they do not establish a universal correction or an absolute benchmark ranking.

## 1 Introduction

TCRdist [1] and its reference implementation tcrdist3 [2] define a widely used distance between paired alpha/beta receptors, and specificity methods are compared by retrieval or by neighbourhood enrichment. Retrieval assumes comparable neighbourhoods, which in high dimension is known to fail: a few *hubs* appear in nearly every neighbour list irrespective of relevance, and *anti-hubs* in none [3]. Correctives that rescale a distance by local neighbourhood density are standard in that literature [4, 5], and have reached biology in single-cell transcriptomics [6].

That receptor neighbourhoods differ in density has previously been established in immunology, a precedent that should be acknowledged before proceeding. Dash et al. [1] previously defined the **NN-distance**, the average TCRdist between a receptor and its nearest neighbours, to characterize repertoire structure, and relate repertoire sampling density to generation probability. tcrdist3 fits a *background-adjusted radius* per meta-clonotype centroid for the same reason [2]. The neighbourhood-enrichment family normalises an observed neighbour count against a generative expectation, ALICE by Pgen and TCRNET against a control repertoire [7], and a recent benchmark places both above tcrdist3 on that task [8]. Frolenkova et al. [9] developed a quantitative framework for receptor embedding geometry and local functional clustering. That work concerns repertoire organization and does not establish the hubness correction evaluated here. We treat the relationship between our correction and that family of methods as an empirical question; Section 3.6 reports the outcome.

We ask how uneven neighbour occurrence affects receptor-distance evaluation, whether local scaling changes same-epitope retrieval, and where that benefit stops. Density adjustment and the scaling estimator have established precedents. Our contribution is their measurement and evaluation in these receptor corpora, including the negative controls and scope boundaries.

## 2 Methods

### Representation and distance

Variable domains were reconstructed for paired alpha and beta clonotypes and placed in the 128-position IMGT frame [10] using ANARCI germline tables [11] for the called V and J genes, with the CDR3 junction placed by the IMGT gap rule. No alignment step was performed. Distances follow TCRdist: CDR1, CDR2 and CDR2.5 at unit weight, CDR3 at weight 3, per-position substitution deficit normalised by the frequency-weighted median deficit of the alphabet and capped at 1. This leaves the substitution alphabet as the only free component. The reimplementation reproduces tcrdist3: Spearman 0.932 over 400,000 pairs, P@1 0.7387 against 0.7380.

### Corpora

VDJdb [12], release of 3 June 2026, capped at 350 clonotypes per epitope: 10,214 clonotypes, 46 epitopes, 23 HLA alleles. 10x dextramers, ITRAP-filtered [13]: 5,722 clonotypes, 43 pMHC, 4 donors. McPAS-TCR [14], after removing its 22.8 % receptor-level overlap with the VDJdb sample: 543 clonotypes. IMMREP22 [15] is evaluated separately under its published train/test protocol. It derives from VDJdb and is not treated as an independent corpus replication.

### Evaluation

A candidate neighbour counts only if it uses a **different V family on both chains**, at fixed donor or fixed HLA allele. This is a diagnostic control, not a use case: without it, retrieval can use germline sharing, and V-gene bias is itself partly HLA-driven and biologically real. The reported quantity is the **macro gain**, precision at depth k divided by the block’s base rate, averaged over blocks of at least 40 queries (30 for McPAS). **Blocks are retained only when 0 < base rate ≤ 0.90**: a saturated block cannot discriminate and a block with no positives cannot be measured, and without the lower bound the estimator is unbounded. The rule removes one dextramer block (base rate 0.9342), six VDJdb blocks at exactly 1.0000 and one at 0.0000, and no McPAS block; both readings are reported. Intervals are a paired bootstrap, 4,000 draws, the same resample indices applied to both methods; because the quantity is a mean **over blocks**, a hierarchical bootstrap that resamples blocks and then queries within them is reported and is the one to read. Labels permuted within donor give the null.

Ties matter here because the distance is a sum of discrete deficits: 10.17 % of dextramer queries have their tenth and eleventh candidates at equal distance under TCRdist against 0.94 % after correction. Ties break stably by row order. Row-order sensitivity was not established independently, and discrete distance ties can change under floating-point rescaling.

### The correction

Hubness is quantified by the occurrence count N_k, summarised by skewness, maximum, and the count of never-retrieved receptors. The correction is local scaling [16], known as a hubness reduction since [4]: ď(i, j) = d(i, j) / sqrt(s_i · s_j), s_i the distance from i to its k-th nearest neighbour. k = 20 was taken as a default and not tuned for the primary correction. Self-distances are excluded when estimating the scale, which is floored at 10^-6 to handle zero distances. **At the rank, s_i cancels**: for a fixed query the ordering is that of d(i, j)/sqrt(s_j), so in retrieval the correction is a density-dependent candidate rescaling. The query-side factor does act wherever scores are compared across queries, as in an AUC. Where a train/test split exists the correction is applied **inductively**, a test point’s s coming from the training set only; and s can equally be read off an external reference set (Section 3.4).

### Study design

Exploratory, without a prospective analysis protocol or a predefined stopping criterion. The macro measure was adopted after inspecting the per-epitope table, because micro precision is degenerate on the dextramer corpus, where one block contributes 1,050 queries at a base rate of 0.9342. The choice reverses rather than confirms conclusions: local scaling scores −0.0015 in micro and +0.2457 in macro on that corpus. The positive retrieval claim therefore depends on the macro endpoint, which was selected after inspection.

### Independent validation corpus and protocol record

IMMREP25 was used once, after the analyses of Sections 3.1 to 3.9 were complete, and for no other calculation in this work. Its 10,000 rows are the cross product of 1,000 distinct paired receptors with the ten epitopes of their HLA; we retain the positive rows only, giving one epitope per receptor, and group comparisons by HLA. An exact paired-CDR3 check found no shared alpha/beta sequence pairs between the 965 retained IMMREP25 receptors and the June 2026 VDJdb human paired records or IMMREP22 training and test records. Adding normalized V and J genes to the identity also yielded no matches. Near-sequence, donor, source-study and representation-pretraining overlap were not assessed. Gene names follow the Adaptive convention and were resolved to IMGT only where a single germline of the ANARCI tables reproduced the CDR1 and CDR2 recorded in the file exactly, a receptor being dropped otherwise. The designs of rounds 37 to 41, including the candidate corrections, the regimes, the decision rule and the outcome to be reported in each case, were recorded in repository protocol files before the corresponding computations, as documented in the analysis records; the files are analysis/PREENREGISTREMENT_tour37.md through PREENREGISTREMENT_tour41.md. Sections 3.1 to 3.8 were exploratory without prospective protocols. Rounds 40 and 41, which support the eight-correction result of Section 3.7 and the whole of Section 3.11, were run after IMMREP25 had been used and are exploratory: their protocols were recorded before computation, but they have not been validated out of sample. These local protocol records are not registrations in an external registry.

## 3 Results

### 3.1 TCRdist over the paired variable domain is hub-dominated

To establish whether retrieval-based evaluation is affected by unequal neighbourhood sizes, we computed the occurrence count N_10_, the number of ten-nearest-neighbour lists a receptor enters, on all three corpora. Occurrence counts were strongly right-skewed and a substantial fraction of each corpus was unreachable (Figure 1). Skewness of N_10_ was +1.2573, +1.3250 and +1.4955 on the dextramer, VDJdb and McPAS-TCR corpora, falling to +0.2472, +0.3090 and +0.1347 after correction, and the number of never-retrieved receptors fell from 178/5,722, 343/10,214 and 13/543 to 13, 22 and 2, that is, from 2.4 to 3.4 % of each corpus to below 0.4 %.

**Figure 1.**
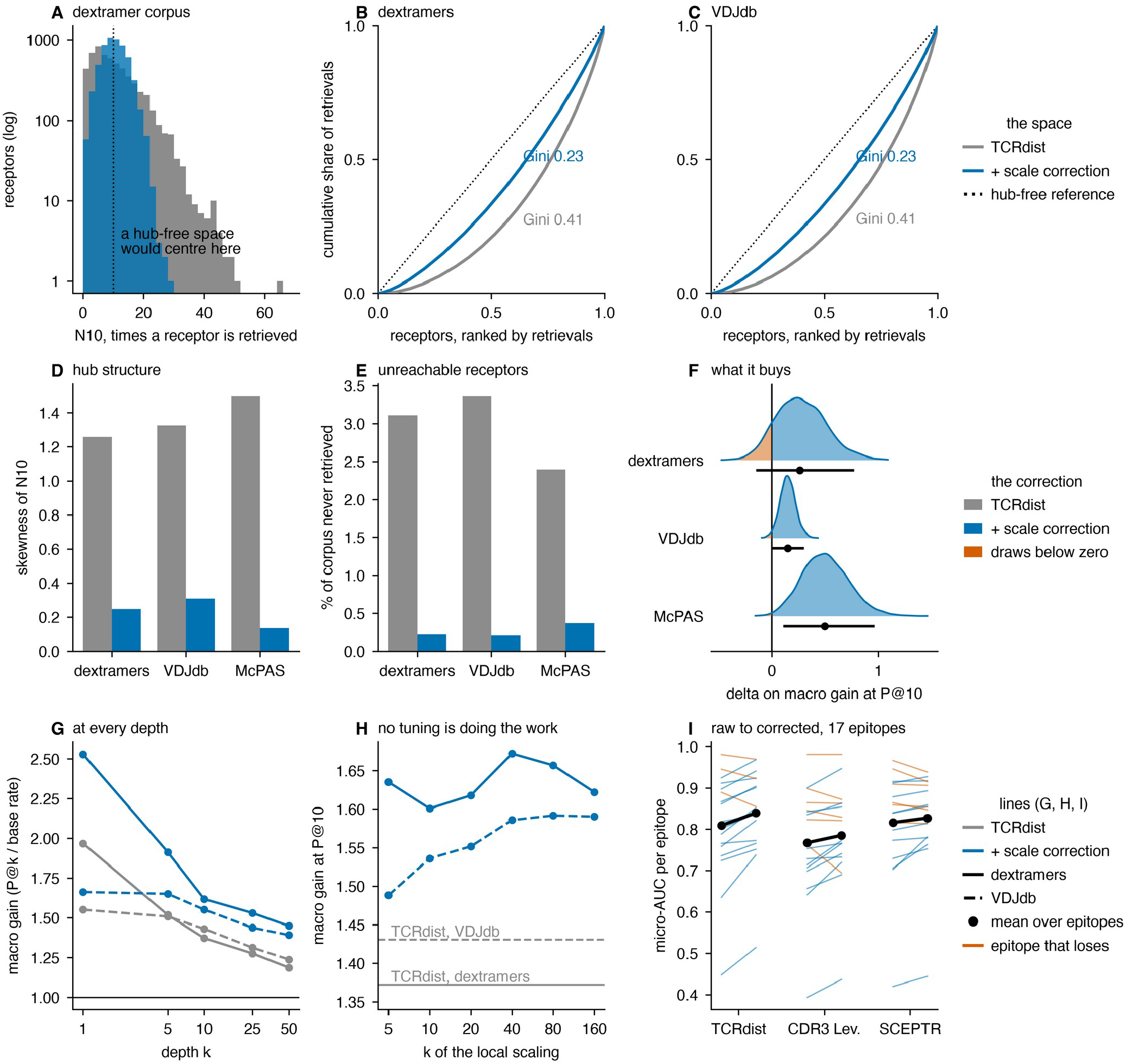
Hub dominance of the space, its removal by local scaling, and the effect on retrieval. (A) Distribution of the occurrence count N_10_ on the dextramer corpus before and after correction, logarithmic ordinate; a hub-free space would centre on N_10_ = 10. (B, C) Concentration of retrievals, on the dextramer corpus and on VDJdb: the cumulative share of all retrievals against the receptors ranked from least to most retrieved, with the diagonal as the hub-free reference and the Gini coefficient beside each curve. (D) Skewness of N_10_ on the three corpora. (E) Percentage of each corpus retrieved by no query. (F) Bootstrap distribution of the delta on macro gain at P@10, four thousand hierarchical draws over blocks; the part of a distribution lying below zero is filled in vermilion, and the bar beneath each density is the 95 % interval with its median. (G) Macro gain against retrieval depth. (H) Macro gain at P@10 against the k of the local scaling, with the uncorrected value as a flat reference; k = 20 was never tuned. (I) Per-epitope micro-AUC before and after correction for three unrelated base representations; black is the mean over epitopes, vermilion an epitope that loses. Panels G and H show the exploratory all-block readings (10 dextramer blocks and 48 VDJdb blocks), whereas panel F and Section 3.2 apply the base-rate exclusion rule.

This is a property of the space rather than of one implementation. The official tcrdist3 implementation and three alternative substitution alphabets showed the same pattern: on the dextramer corpus the skewness of N_10_ before and after correction was +1.26 → +0.25 for BLOSUM62, +1.26 → +0.26 for BLOSUM45, +1.30 → +0.31 for PAM70, +1.21 → +0.25 for an identity matrix and +1.19 → +0.25 for official tcrdist3, with the query-resampling gain intervals used in that analysis excluding zero in each case. These are not the hierarchical block intervals used for the primary result (per-distance gains, intervals and skewness in analysis/tables/tour20_generique.csv).

### 3.2 Local scaling reduces hub structure and improves same-epitope retrieval

We then rescaled each pairwise distance by the geometric mean of the two receptors’ local neighbourhood scales (local scaling, k = 20, untuned). Same-epitope retrieval improved on all three corpora, and on VDJdb the block-level interval excluded zero.

| corpus | blocks | TCRdist | corrected | delta, blocks resampled | blocks improved |
| --- | --- | --- | --- | --- | --- |
| VDJdb | 41 | 1.5284 | 1.6702 | +0.1418 [+0.0079; +0.3016] | 26/41 |
| McPAS-TCR | 7 | 1.0253 | 1.5211 | +0.4957 [+0.1068; +0.9634] | 6/7 |
| dextramers | 9 | 1.4131 | 1.6865 | +0.2734 [-0.1469; +0.7750] | 6/9 |

Labels permuted within donor placed the observed deltas at z = +10.9, +10.4 and +5.2 from the null, which tests whether the distance is related to the label at all rather than whether the effect is of the size reported. Splitting the VDJdb epitopes into two disjoint halves gave +0.1383 and +0.1095, so the gain is not carried by a small set of epitopes, and every k from 5 to 160 left it above baseline.

The result is carried by VDJdb and not by the dextramer corpus. On the dextramers the block-level interval contains zero, and the cause is concentration rather than sample size: two blocks with base rates of 0.0405 and 0.0091 account for 113 % of the unfiltered delta while four of the ten blocks are negative, and a macro gain whose denominator is 0.0091 amplifies a small number of successes. VDJdb is more favourable on every count, and the dextramer figures are reported for completeness rather than as support.

### 3.3 The gain transfers to an externally fixed benchmark and to unrelated representations

On IMMREP22 the splits, the inductive protocol and the micro-AUC metric were fixed by prior work [15], the classifier is the weakest available, the distance to the nearest training positive, and the presence of the correction is the only variable. Receptors were dropped for want of an ANARCI-resolvable gene at a rate of 0.34 % of the training data (50 of 14,701) and 0.38 % of the test data (14 of 3,675).

Mean micro-AUC over the seventeen epitopes rose from 0.8091 to **0.8394**, improving 14 of 17 (paired Wilcoxon p = 0.0026), with the largest gains on the hardest epitopes. Because an AUC compares scores across test points, the two factors of the formula separate: the candidate side alone gave 0.8171 and the query side alone 0.8334. The larger part of the benchmark gain is therefore a per-test-point score normalisation rather than the candidate-side rescaling that carries the retrieval result of Section 3.2. We report the paired delta only: the published entries each bring their own data preparation, and a bootstrap over epitopes does not support a claim about absolute placement.

The correction is not specific to TCRdist. Applied unchanged on the same benchmark, it raised a CDR3 Levenshtein distance from 0.7675 to 0.7853 (+0.0179, 11 of 17, p = 0.056) and the contrastively trained representation SCEPTR [17] from 0.8162 to 0.8271 (+0.0109, 11 of 17, p = 0.19); 36 of the 51 (representation × epitope) pairs improved, with a median delta of +0.0161 (Figure 1I). Only the TCRdist row is significant on its own, so the generality of the post-processing reading rests on the direction being the same across a hand-built distance, an edit distance and a neural embedding, not on any one of them. The SCEPTR space is itself hub-dominated (skewness of N_10_ +1.03 on the dextramers and +1.25 on VDJdb), so hub dominance is not an artefact of hand-built distances.

### 3.4 The scale can be read from an external reference set, including a synthetic repertoire

Read off an external reference set rather than the corpus under evaluation, the correction becomes fully inductive and applicable to a single receptor pair. Using 14,963 paired memory-CD8 receptors carrying none of our labels, it gave +0.1181 [+0.0092; +0.2397] on VDJdb and 0.8457 on IMMREP22 (16 of 17, p = 7.6 × 10^-5^), that is, smaller than the corpus-derived scale on the dextramers, larger on the benchmark, and less variable. On McPAS-TCR the same external scale gave +0.0039 [−0.1951; +0.2330], no detectable effect. IMMREP22 draws its negatives from that same control set, so each epitope’s own training and test receptors were removed from the reference before use, 357 on average.

The retrieval results in this section use the exploratory all-block reading; the corresponding filtered VDJdb external-scale delta is +0.1382 in Section 3.6.

Twenty-six reference sets were then compared (Figure 3A to 3C). In the tested reference sets, two hundred and fifty receptors were sufficient, as was each single-donor reference and a synthetic repertoire drawn from the V(D)J recombination model before thymic selection [18], on the dextramers and on the benchmark alike, so a synthetic scale reference can support this correction in the tested settings. Two reference sets constructed to be adversarial did not fail: re-pairing the alpha and beta chains at random, and using a small specificity-enriched set, both performed about as well as the intended reference. The reference set is therefore a scale reference and not a null model, and the objective differs from a background-adjusted radius [2], since that construction requires an unenriched repertoire whereas this one requires only some sample of TCR-like sequences.

**Figure 2.**
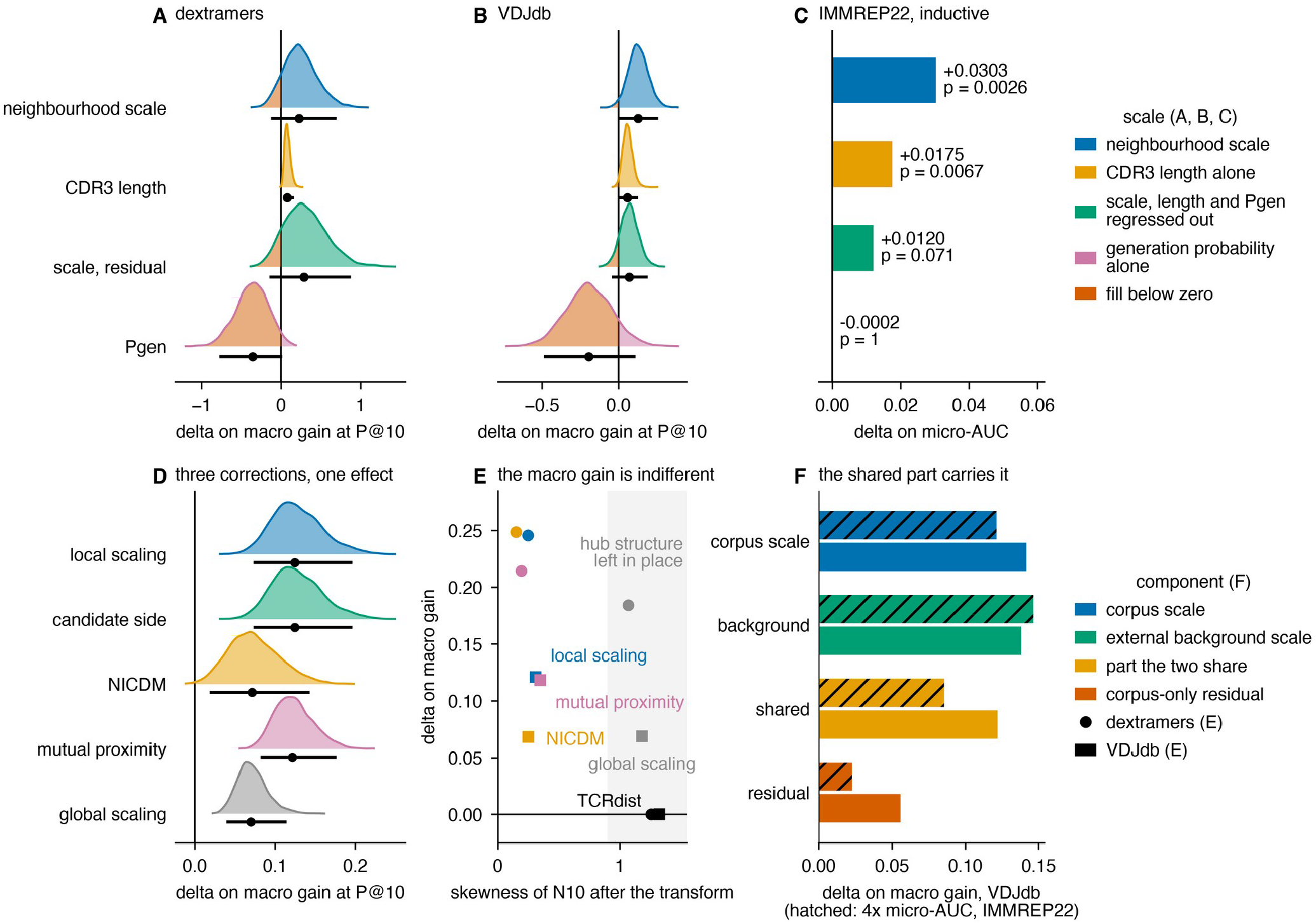
Composition of the scale, and alternatives that match it. (A, B) Bootstrap distributions of the delta on macro gain at P@10 for four scales on the dextramer corpus and on VDJdb: the neighbourhood scale, CDR3 length alone, the neighbourhood scale with length and generation probability regressed out, and generation probability alone. (C) The same four on the IMMREP22 mean micro-AUC, inductive, with the Wilcoxon p beside each bar. (D) Four neighbouring corrections and one global-scale control on VDJdb. These draws follow the resampling scheme of round 24, which resamples queries within fixed blocks rather than resampling the blocks, and are therefore narrower than every other interval reported here; the difference lies in the scheme, not in the estimate. (E) Delta on macro gain against the skewness of N_10_ after the transform, circles for dextramers and squares for VDJdb; global scaling matches the macro measure while leaving the hub structure in place (Section 3.8). (F) The corpus scale, the external background scale, the part the two share, and the corpus-only residual; the IMMREP22 bars are micro-AUC deltas and are drawn at four times the scale of the VDJdb macro-gain bars so that both fit one axis. Panels A and B use the exploratory all-block readings, as in the table in Section 3.5.

**Figure 3.**
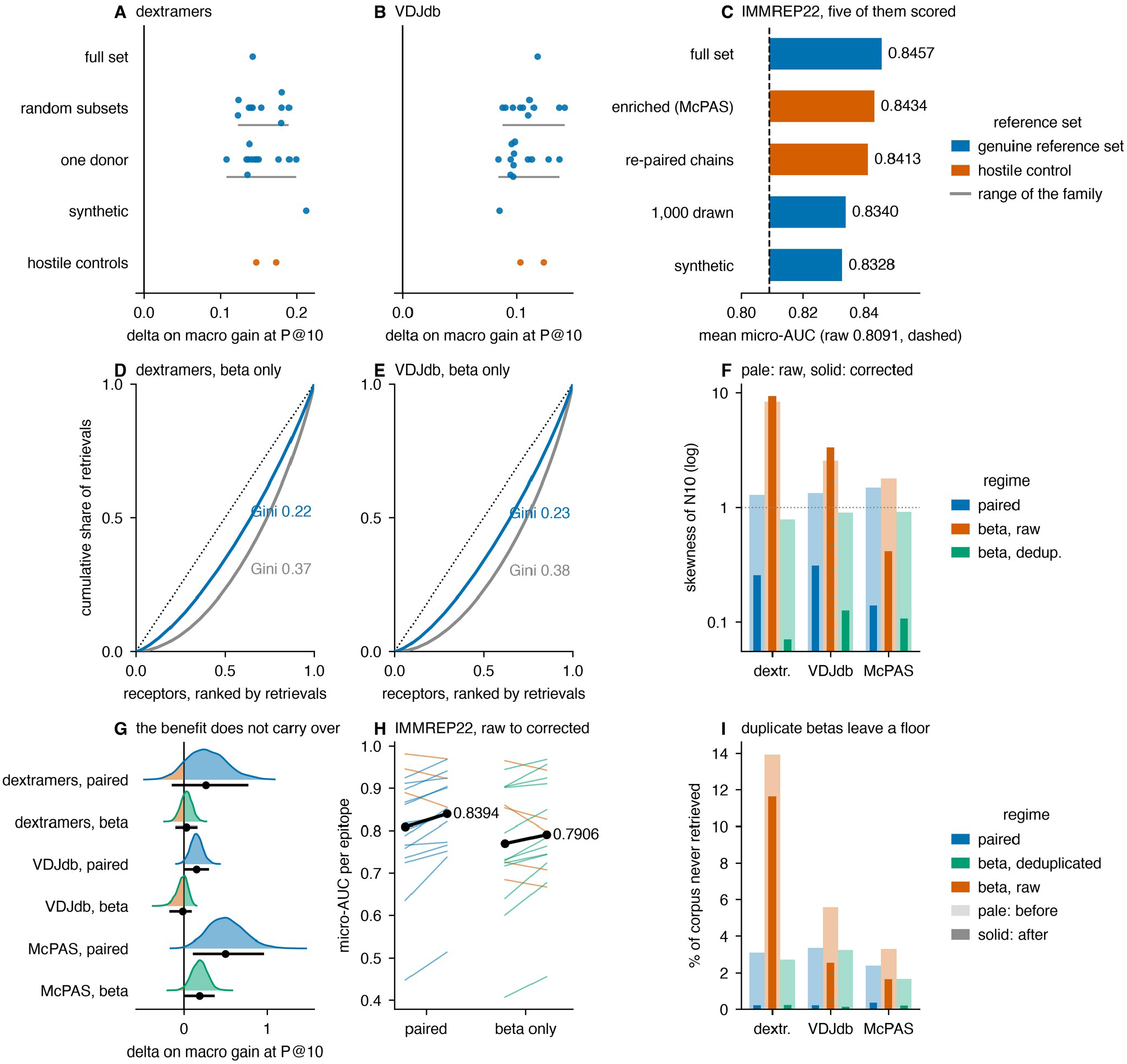
Scope of the correction: choice of reference set, and the single-chain regime. (A, B) Delta on macro gain at P@10 for twenty-six reference sets, on the dextramer corpus and on VDJdb, one point per set, grouped by family: the full control set, random subsets from 250 to 5,000 receptors, each of twelve donors taken alone, a synthetic repertoire drawn before thymic selection, and the two adversarial reference sets in vermilion. The grey rule under a family spans its range. (C) The five reference sets also scored on IMMREP22, against the uncorrected mean micro-AUC (dashed); the axis is truncated. (D, E) Concentration of retrievals on the deduplicated beta chain alone, read as in Figure 1B. (F) Skewness of N_10_ before (pale) and after (solid) correction, for the paired domain, the raw beta chain and the deduplicated beta chain; the raw beta arm is shown to make the duplicate confound visible and is not interpretable as a result. (G) Bootstrap distributions of the delta on macro gain, paired against deduplicated beta. (H) Per-epitope micro-AUC on IMMREP22, paired against beta; the beta arm there is not deduplicated. (I) Percentage of each corpus retrieved by no query, before (pale) and after (solid) correction, in the three regimes: duplicate betas leave a floor that no rescaling can lift. Panels A and B use all blocks meeting the query-count threshold, without the base-rate exclusion rule.

### 3.5 The scale is not reducible to CDR3 length or generation probability

We next replaced the neighbourhood scale s by quantities that consult no neighbourhood at all (Figure 2A to 2C).

| scale | VDJdb, delta | IMMREP22, delta |
| --- | --- | --- |
| neighbourhood scale $s$ | +0.1211 [+0.0054; +0.2595] | +0.0303 (14/17, $p = 0.0026$) |
| CDR3 length alone | +0.0585 [+0.0041; +0.1291] | +0.0175 (12/17, $p = 0.0067$) |
| $s$ with length and Pgen regressed out | +0.0653 [−0.0440; +0.1923] | +0.0120 (10/17, $p = 0.071$) |
| generation probability alone | −0.1959 | −0.0002 (9/17, $p = 1$) |

The VDJdb column of this table is the all-block reading (48 blocks), which is the state in which round 31 was run; the 41-block reading under the block rule is the one given in Section 3.2. Both readings are reported for every round in analysis/logs/tour32.log.

Two results follow, one restricting the claim and one supporting it. Restricting it: CDR3 length alone, with no distance computed, recovers between a third and three-fifths of the benefit (0.0814 of 0.2457 on the dextramers, 0.0585 of 0.1211 on VDJdb, 0.0175 of 0.0303 on IMMREP22), and the neighbourhood signal remaining once length and generation probability are regressed out is not significant on its own on either VDJdb or the benchmark. Supporting it: generation probability used as the scale does not merely fail to reproduce the effect, it degrades retrieval, and log s is explained by generation probability and length with an R^2^ of only 0.030 on VDJdb and 0.140 on the dextramers. These controls do not support explaining the gain by generation probability alone. On VDJdb, length-only scaling gives +0.0585, with an interval excluding zero. The residual scale gives +0.0653, with an interval containing zero. These effects are not an additive decomposition of the full-scale retrieval gain.

In the earlier exploratory analysis, two further hubness reductions had positive deltas on VDJdb, NICDM at +0.0681 [+0.0189; +0.1434] and mutual proximity at +0.1183 [+0.0820; +0.1766], with query-resampling intervals in that analysis (Figure 2D). These intervals do not replace hierarchical block inference or establish generality across correction methods.

### 3.6 The scale is not a restatement of neighbourhood enrichment

Rescaling a distance by local crowding and testing a neighbourhood for enrichment against a baseline could be two expressions of a single operation. Two tests, with different nulls, indicate that they are not.

The first is the result of Section 3.5. Generation probability is the null of ALICE, and used as the scale it does not merely fail to reproduce the effect but degrades retrieval.

The second addresses TCRNET, whose null is a control repertoire rather than a generative model. Enrichment is a ratio of observed to expected local density, and both of its ingredients carried the retrieval gain on their own: the corpus scale at +0.1418 on VDJdb and the external background scale at +0.1382. Their ratio carried none of it: +0.0076 [−0.1464; +0.1802] on VDJdb and −0.0174 on IMMREP22 (4 of 17 epitopes, p = 0.027), significantly worse than leaving the distance unchanged.

This is not a variance artefact. The ratio retains 95 % of the variance of the log corpus scale on VDJdb and 88 % on the dextramers, so signal remained available to measure. Nor is it an artefact of fixing the exponent at one. Fitting log s on the log background scale by least squares gave b = 0.750 on VDJdb, b = 1.211 on the dextramers and a median of 0.475 across the benchmark’s epitopes, and with the fitted coefficient the corpus-specific residual still carried nothing: +0.0559 [−0.0796; +0.2147] on VDJdb, +0.0057 (p = 0.35) on IMMREP22, and on the dextramers it was significantly harmful at −0.3247 [−0.6126; −0.0353]. The fitted common component carried the gain on all three (Figure 2F): +0.1219 [+0.0222; +0.2324] on VDJdb, +0.1990 [+0.0687; +0.3925] on the dextramers and +0.0214 on IMMREP22, with an interval excluding zero even on the corpus where the interval of the raw corpus scale did not.

The interpretation is that the active quantity is the part of the local scale on which a labelled corpus and an unrelated repertoire agree, and that the corpus-specific residual, which holds most of the variance, carries none of the benefit. Neighbourhood enrichment isolates that residual. This also accounts for the behaviour of the adversarial reference sets of Section 3.4, because the tested reference sets retained a useful scale component. Generalization to arbitrary references is not established.

Three caveats narrow this conclusion. First, the fitted common component is by construction a monotone function of the background scale, so it is not independent evidence that the background scale works, which Section 3.4 had already established. Second, a first pass expressed the same quantities as counts within a fixed radius rather than as radii and is not interpretable: its own control shows the background density failing as a count (−0.1580 on IMMREP22) where it succeeds as a radius (+0.0366), so that pass measured the functional form rather than the concept; it is retained in the repository with the amendment dated. Third, this is an enrichment statistic in the geometry used here, paired and over the whole variable domain, and not TCRNET itself, which operates on the beta chain alone at Hamming distance one. These controls distinguish the tested radius-based transformations. They do not establish the mechanism of ALICE or TCRNET, or replace a head-to-head comparison on their own task.

### 3.7 The diagnostic survives on a single chain; the retrieval benefit does not

Most repertoire data are bulk beta-chain sequences, and the neighbourhood-enrichment methods operate in that regime, so whether the preceding results hold outside the paired variable domain determines how far they reach. Recomputing the distance on the beta chain alone divides the answer in two. In the beta-only arm, germline control excludes candidates sharing the query’s TRBV family; the paired arm excludes candidates sharing either its TRAV or its TRBV family.

One confound has to be cleared first, and it is substantial. A corpus deduplicated at the level of the paired clonotype is not deduplicated at the level of the beta chain: 37.3 % of the dextramer corpus and 16.2 % of VDJdb share a beta with another receptor, and the largest group holds 187 identical betas. In single-chain geometry those receptors sit at distance zero from one another and so generate hubs by construction, which is why the raw beta skewness of N_10_ reads +8.34 on the dextramers where the paired reads +1.29. The duplication is also not label-neutral: 20.1 % of the dextramer corpus and 3.6 % of VDJdb carry a beta tied to more than one epitope, which no single-chain method can separate. That is a property of the data rather than of any method. The interpretable arm removes ambiguous betas and then deduplicates; the confounded arm is retained in the repository.

The diagnostic holds (Figure 3D to 3I). On the deduplicated beta chain, skewness of N_10_ fell from +0.7857, +0.8993 and +0.9133 to +0.0706, +0.1264 and +0.1074 across the three corpora, and never-retrieved receptors fell from 105/3,851, 293/8,994 and 8/480 to 9, 12 and 1. Before correction the beta space is in fact less hub-dominated than the paired one, which is what the hubness literature predicts of the lower-dimensional of two spaces [3].

The retrieval benefit does not hold. The delta on macro gain was +0.0285 [−0.1044; +0.1615] on the dextramers and −0.0223 [−0.1709; +0.0926] on VDJdb, and was positive only on the smallest corpus, +0.1829 [+0.0076; +0.3695] on McPAS-TCR. On IMMREP22 the beta arm gave +0.0208 (12 of 17, p = 0.051), and that arm is not deduplicated: 8.3 % of test betas on average and 16.7 % at worst are already present in the training set, so a borderline figure is if anything optimistic.

This was established with one correction, and the corrections do not all behave alike (Sections 3.9 and 3.10), so we repeated it with eight: local scaling at k = 5, 20, 50 and 100, NICDM, mutual proximity, the global scale and the external-background scale. The deduplicated beta arm gives sixteen cells on the two principal corpora and every interval contains zero; the largest are +0.1338 [-0.0568; +0.3847] for mutual proximity on the dextramers and +0.0364 [-0.0605; +0.1204] for local scaling at k = 100 on VDJdb. Two cells on McPAS-TCR, the corpus with six blocks, have intervals excluding zero and neither survives the Holm correction over the eight candidates that was declared in advance. Local scaling reproduced the three figures above exactly, which is the internal control on this arm. The boundary persisted across the tested estimators; this does not establish a universal property of every single-chain representation.

Removing hub structure and improving retrieval are therefore not the same operation. On a single chain the correction still removes the hubs and no longer improves retrieval. The result of Section 3.2 accordingly carries a scope condition: it belongs to the paired variable domain, the higher-dimensional of the two spaces, and it is there that correcting pays.

### 3.8 Macro gain alone does not certify a correction

Dividing by each receptor’s mean distance to the whole corpus is a global scale: one number per receptor, no neighbour consulted, and no hubness rationale. In the all-block sensitivity analysis, it outperformed TCRdist on both large corpora with intervals excluding zero, +0.1842 on the dextramers and +0.0687 on VDJdb (Figure 2E), and local scaling did not outperform it on this measure, the paired block-level interval containing zero on all three corpora (+0.0616, +0.0524 and +0.3477). What separates the two is the hub structure, and there the separation is not marginal: local scaling took the occurrence-count skewness from +1.33 to +0.31 on VDJdb, whereas the global scale left it at +1.18.

The inductive benchmark does separate them. On IMMREP22, seventeen epitopes under a third party’s protocol and splits, TCRdist scored 0.8091, local scaling 0.8394 and the global scale 0.8215. Local scaling outperformed the global scale with a paired Wilcoxon p = 0.0129, winning on thirteen of the seventeen epitopes. The macro-gain comparison did not establish a significant difference between the two scales, whereas the inductive AUC comparison did. The two evaluation measures therefore support different conclusions.

Macro gain alone consequently cannot carry the argument, not because it rewards a collapsed space but because it is indifferent to the mechanism. The argument requires separate evidence from retrieval, neighbour occurrence and the inductive benchmark AUC.

Local and global hubness corrections have established precedents [4, 5]. Our mean-distance control is defined explicitly above; it should not be equated with every method called global scaling in that literature. The comparison here concerns the measured behavior of this control in receptor space.

An earlier version of this analysis reported a delta of +2.2028 for this control and read it as a degenerate transform collapsing the corpus. That figure came from a defective computation, since corrected; see ERRATA.md and round 36.The conclusion of the section is unchanged and now rests on the corrected control rather than on an artefact.

### 3.9 Reducing hub structure and improving retrieval are different objectives

For a fixed query the scale of that query is constant along its row, so the order induced by d(i,j)/sqrt(s_i s_j) is the order induced by d(i,j)/sqrt(s_j). In retrieval the correction is therefore not a symmetric rescaling of the two receptors but a one-sided penalty on the candidate. We verified this rather than asserting it: for NICDM and the global scale, on three corpora in two regimes, the macro gain of the full transform and of its candidate half agree to within 1e-9 in all twelve comparisons, local scaling itself having been checked the same way earlier. The query factor does not cancel on the inductive benchmark, where scores rather than ranks are compared, and Section 3.3 reports that it carries the larger part of the gain there. The two statements are consistent and apply to different measures.

If the benefit arose from suppressing hubs as such, then penalizing the occurrence count itself should be its strongest form. It is its worst. In unrestricted within-group retrieval, multiplying each distance by the square root of one plus the candidate’s occurrence count lowers the macro gain by 2.6120 on the dextramers [-5.1619; -0.5503] and by 1.7734 on VDJdb [-2.4850; -1.1243], both p = 0.0003, improving 2 of 9 and 6 of 42 blocks. This is the largest effect reported in this work and it is negative. Receptors that enter many neighbour lists are frequently the correct answer, and a penalty that targets that property directly removes signal. What local scaling penalizes is local density, not hub status; the two proxies point in opposite directions, and only the first is useful. The observed occurrence imbalance is a useful diagnostic. Its reduction alone does not establish improved specificity retrieval.

### 3.10 The correction is confirmed out of sample under germline control, and only there

We recorded a prospective test of universality: four candidate corrections, local scaling, NICDM, mutual proximity and the global scale, across three corpora and three regimes, with a correction being called universal only if its interval excluded zero from above in all nine cells with a Holm-corrected p below 0.05. None did; the best passed one cell of nine. Across the open regime, the regime that corresponds to the use case, rounds 37 to 39 evaluated thirty-three candidate cells. One interval excludes zero from above, +0.0334 on VDJdb for the global-scale control, and it does not survive the multiplicity correction declared in advance (p = 0.086).

We then validated the published correction on IMMREP25, a corpus used for no other calculation in this work and read once. Its receptors carry Adaptive gene names, which we resolved to IMGT only where a single germline reproduced the observed CDR1 and CDR2 exactly; 35 of 1,000 receptors (3.5 %) could not be resolved and were dropped, leaving 965 receptors across 20 blocks. Under germline control, local scaling with the untuned k = 20 of the rest of this work gives +0.1185 in macro gain, block-level interval [+0.0040; +0.2477], p = 0.041, improving 13 of 20 blocks. Two candidates found by searching the method space after seeing the results of round 37 were tested in the same pass and neither was confirmed: local scaling at k = 100, which had been the strongest variant on VDJdb, gives +0.0990 [-0.0085; +0.2069] with a point estimate below that of k = 20, and the external-background scale gives +0.1659 [-0.0155; +0.4051]. The apparent superiority of the larger k was specific to the corpus on which it was found. In the open regime on this corpus all three readings lie between +0.0093 and +0.0233 with intervals containing zero.

This held-out result supports the correction in the germline-controlled setting. The lower interval bound is +0.0040 and the result is marginal. Before gene-resolution exclusions, IMMREP25 contains fifty positives per epitope. This constructed balance differs from an unselected natural repertoire, so transfer to such a repertoire is not established by it. Having been used once, it is no longer available as an independent validation set for this project.

### 3.11 The corrected distance is not a metric, and that is not why clustering behaves differently

Local scaling can violate the triangle inequality. We tested whether repairing that violation restores the clustering behavior of the uncorrected distance.

The violation is real. Within donors, local scaling breaches the triangle inequality by up to 9.0917 on the dextramers and 4.9761 on VDJdb, on an average 0.25% and 2.14% of triples per evaluated group. Checks were exact in smaller groups and sampled in larger groups. The uncorrected numerical violations were at most 2.3e-5, consistent with floating-point error. We repaired it in two ways and reclustered. Adding a common constant to off-diagonal distances leaves their order unchanged and, in exact arithmetic, shifts all average-linkage merge distances equally. The implementation showed small numerical deviations at some fine partitions on VDJdb; we therefore do not claim exact identity of every computed partition. For sampled groups, the constant addresses the observed maximum violation rather than certifying every triangle. The metric closure, computed by shortest paths within each donor, does change the ranks, and changes almost nothing: adjusted Rand 0.0391 against 0.0367 on the dextramers and 0.0784 against 0.0777 on VDJdb, at a cost of 0.0108 and 0.0074 in macro gain. Non-metricity is therefore not what separates the two distances under average linkage.

What separates them is the granularity at which each works. The correction moves the useful number of clusters downwards. On VDJdb it exceeds TCRdist on adjusted mutual information at all five cluster counts tested, 0.2351 against 0.0033 at the coarsest, and on adjusted Rand at three of five; on McPAS-TCR it leads on three of five for both indices; on the dextramers it leads on two of five and one of five. Each distance peaks at a different granularity, TCRdist at four to sixteen times the number of epitopes and local scaling at one to two times. A comparison made at a single fixed number of clusters therefore reports the choice of that number as much as the distance, and the pre-registered rule used here read both methods at the count where TCRdist peaks, which is the reading least favourable to the correction. Clustering does not simply fail to improve; it improves on the coarse partitions and degrades on the fine ones, and which of those a benchmark reports decides the answer.

Rounds 40 and 41 are exploratory. They followed the validation of Section 3.10, which consumed the only corpus this work held in reserve, so nothing in this section or in the eight-correction result of Section 3.7 has been confirmed out of sample.

## 4 Discussion

Receptor-distance benchmarks contain unequal neighbour occurrence, and changing the local scale can change both retrieval and score calibration. This is the best supported of the results reported here: it is a direct measurement rather than an inference, and it is inexpensive to reproduce on any labelled repertoire. The correction that follows from it improves retrieval, but within boundaries that are as informative as the gain itself, and it does not work by the route its name suggests: penalizing hub status directly is strongly harmful (Section 3.9), so hubness is what these benchmarks partly measure rather than a quantity to be minimized.

### Recommendations

Two quantities should accompany any receptor-distance benchmark that reports precision at a fixed depth: the skewness of the occurrence count, and the fraction of the corpus retrieved by no query. Both are inexpensive to compute. A reported retrieval improvement should further be accompanied by the same method evaluated with and without a scale correction, since the correction applied here raised the mean micro-AUC of all three base representations tested on an externally fixed benchmark. The comparison should report the retrieval and hubness outcomes separately: improvement in one does not establish the mechanism of the other.

#### Limitations

ordered by how much they restrict the claim.

#### The evaluation measure does not separate the two mechanisms

(Section 3.8) and was chosen post hoc. Sections 3.1 to 3.8 were exploratory without prospective protocols; the universality test, the search over further candidates and the IMMREP25 validation were (Section 2).

#### CDR3 length alone recovers between a third and three-fifths of the effect

and the neighbourhood residual is not significant on its own on VDJdb or on the benchmark.

#### The larger part of the benchmark gain is a per-test-point score normalisation

(Section 3.3), not the candidate-side rescaling. In retrieval the query factor cancels at rank altogether (Section 3.9), so the two measures are carried by different halves of the same transform: on IMMREP22 the candidate side alone gives 0.8171 and the query side alone 0.8334, against 0.8394 for the two together.

#### The retrieval benefit is confined to the paired variable domain

(Section 3.7). On the beta chain alone none of eight corrections yields an interval excluding zero in the deduplicated beta-only germline-controlled arm of either principal corpus, while the hubness diagnostic survives intact, and most available repertoire data are single-chain. Measured with eight corrections on three corpora, it is no longer the weakest supported of these limitations.

#### The dextramer corpus is unreliable here

(Section 3.2): two low-base-rate blocks dominate its delta and its block-level interval contains zero. It is also the corpus where epitope and donor are confounded and where the raw data are heavily filtered [13].

#### The germline-controlled regime is a diagnostic, not a use case

Outside it, no retrieval gain survived the declared multiplicity corrections across four corpora and thirty-three candidate cells (Section 3.10). This is a limit on the evidence for a gain, not a demonstration of zero effect. In that regime the V gene already carries most of the retrieval, the TCRdist macro gain on the two principal corpora rising from 1.4131 and 1.5284 under germline control to 4.7389 and 5.0022 without it, which leaves little for any correction to recover.

#### Clustering changes character rather than simply improving

(Section 3.11). The correction leads on coarse partitions and loses on fine ones, and a comparison at one fixed number of clusters reports that choice as much as the distance. The corrected distance is also not a metric, which restricts its use in the tools the field clusters with, but repairing the violation moves adjusted Rand by less than 0.003, so the two facts are independent rather than one explaining the other.

#### The comparison to other methods is partial

SCEPTR was scored; GLIPH2, TCRMatch, clusTCR and GIANA were not. Radius-based controls related to neighbourhood enrichment were evaluated (Section 3.6); ALICE and TCRNET themselves were not scored on their own task.

#### Outlook

Whether a distance-level scale correction and a neighbourhood-enrichment test are two expressions of a single operation is an empirical question, and Section 3.6 answers it against that identification: under two distinct nulls, a generative model and a control repertoire, the active quantity is not the excess of the corpus over a background. The comparison that remains is narrower, namely scoring ALICE and TCRNET on their own task in their own single-chain geometry. Section 3.7 sets the terms on which such a comparison should be read, since on a single chain it is the diagnostic, and not the retrieval benefit, that survives.

## Data and code availability

The analysis repository is https://github.com/KilianMaire/tcr-hubness. It contains scripts, analysis logs, tables and protocol records. The original corpora are obtained from their providers: ITRAP-filtered dextramers [13], VDJdb [12], McPAS-TCR [14] and IMMREP22 [15]. IMMREP25 was obtained as immrep2025_for_release.tsv with the provider’s dataset and methods document. Provider access and reuse conditions apply.

## Ethics

This study reanalyses existing datasets and reports no new participant recruitment or experiments. Dataset-specific ethical approvals and reuse conditions remain those of the original studies.

## Notes

https://github.com/KilianMaire/tcr-hubness

### Competing Interest Statement

The authors have declared no competing interest.

